# The Earlier You Know, the Smoother You Act

**DOI:** 10.1101/2025.11.08.687371

**Authors:** Abir Chowdhury, Heiko Maurer, Alap Kshirsagar, Kai Ploeger, Jan Peters, Hermann Müller, Lisa Katharina Maurer

## Abstract

Intercepting a moving object with our hands requires that the moving object and the hand come into contact at some point along the object’s trajectory. Hand movement must, therefore, be planned in advance so that the hand reaches a chosen spatial location along this trajectory at the correct moment in time. To enable such anticipatory behavior, the object’s trajectory must be predicted based on sensory information collected before movement planning is concluded. Early completion of planning would allow for a smooth and economical movement; however, it also requires sufficiently accurate information about the object’s movement to be available earlier. We investigate interception in the context of a complex, naturalistic motor task, toss juggling, to understand the extent to which proprioceptive and tactile information influence anticipatory movements of the catching hand. In our study, we compared solo juggling, where the efferent, tactile and proprioceptive information from the throwing hand is available during catch planning, with dyadic juggling, a functionally “deafferented” condition where no such information is available because the ball has been thrown by another person. Our results indicate that when internal and tactile information from the toss is absent, anticipatory behavior is greatly reduced. Experienced jugglers seem to rely more strongly on internal information than vision, allowing earlier planning and lower target uncertainty. In the dyadic condition, when internal information is absent, trajectory planning is delayed, leading to greater target uncertainty.

## 1 Introduction

Interception is a fundamental component of everyday motor tasks, particularly in dynamic activities such as hitting a moving ball in sports or, in more static contexts, reaching for a glass on a table. In the context of a dynamic activity, it can be described as the process of coordinating spatial and temporal parameters to arrive at a specific location at the precise moment another object reaches that point, thereby enabling physical interaction or contact with the object. A central theory in motor control proposes that accurate movement relies on the brain forming internal models that predict the sensory consequences of motor commands (Wolpert et al. 1995, 1998; Kawato 1999). In ball-catching tasks, the brain uses sensory information about the ball movement to generate predictive estimates of the ball’s future trajectory, enabling anticipatory motor planning (Hayhoe and Ballard 2005; Diaz et al. 2013; de Lussanet et al. 2004). At a certain point in the movement, the internal model reaches sufficient accuracy, allowing for effective anticipation of the ball’s path to guide interception. Finally, toward the final phase of the movement, only minor corrective adjustments are made (López-Moliner et al. 2010).

Toss juggling is an extreme example of such motor activity that demands rhythmic catching and throwing of objects under spatial and temporal constraints (Beek and Turvey 1992). The dense time schedule of juggling requires the juggler to initiate the final approach toward the intended interception position while simultaneously coordinating the throw of the ball already in hand. Unlike discrete object-catching, where the hand often comes to a stop after interception, juggling involves continuous motion, each catch serves as a preparation for the next throw. For instance, in a cascade pattern, this seamless coordination is crucial for maintaining a stable juggling rhythm, as the quality of each throw directly affects the ease of the following catch of the same object by the other hand. After all, the difficulty in juggling increases due to the simultaneous control of multiple objects. However, balls are not thrown to random locations; rather, the trajectory of each throw is structured and predictable. As a result, each ball does not need to be visually tracked or caught independently. Skilled jugglers capitalize on this predictability by relying on the accuracy of their throws and internal predictions of where each ball will travel. Remarkably, up to five balls can be juggled while blindfolded, underscoring the reliance on internal models rather than continuous visual tracking (Botvinick-Greenhouse and Shinbrot 2020). The degree of accuracy required in these throws reflects the inherent instability, and thus the difficulty, of a given juggling pattern. Experienced jugglers further optimize performance by fixating near the apex of the juggling arc, rather than following each ball individually (Dessing et al. 2012, 2009). This gaze strategy, along with multisensory integration from proprioceptive information about the own movements as well as tactile and visual information about the ball’s behavior and a finely tuned internal model, enables them to maintain control and rhythm in an otherwise highly unstable and dynamic motor task. This capacity to integrate multiple actions into a continuous process distinguishes experienced jugglers from novices, who often exhibit disjointed and reactive movements. Ultimately, juggling exemplifies a highly efficient form of motor control in which multiple actions are anticipated and executed with precision to sustain uninterrupted motion.

One way to analyze the different components of coordinated juggling and their interrelations is to use the minimum jerk trajectory-based approach proposed by Slupinski et al. (2018). With this approach the planning process of catching can be computed from the shape of the hand-movement trajectory. The fluidity in the catching movement is achieved through a combination of feedforward control, which enables prediction-based adjustments before the ball arrives, and feedback control, which allows for real-time corrections when deviations occur. This characteristic is manifested in a bell-shaped velocity profile typical for goal-directed reaches. The term “minimum jerk trajectory” refers to the smoothest possible movement between two points over a given time: it minimizes the integral of squared jerk (the rate of change of acceleration) (Flash and Hogan 1985). Because such profiles are taken as signatures of predictive, feedforward execution, the onset of the smooth-approximation phase marks the beginning of feedforward-driven (predictive) control (Shadmehr and Wise 2004). An earlier onset suggests an earlier start of predictive planning, whereas a later onset indicates continued reliance on sensory feedback before the movement settles into a smooth trajectory. By relating the timing of this smooth-approximation onset to the release time of the toss from the other hand, we can estimate when the catching hand begins to follow a smooth path toward the ball, serving as a proxy for the completion of planning for the initial ballistic approach to the catch.

Previous work has identified the onset of smooth, goal-directed movements in discrete interception tasks using self vs non-self throws (Slupinski et al. 2018; Hagenfeld et al. 2022). In the present study, we investigate movement planning in a more complex interception task, toss juggling, where the hand transitions fluidly between throw and catch phases. From the hand’s perspective, the process of catching a ball during juggling involves continuously adjusting its movement to get closer to where the ball is expected to be. At each moment, we can calculate the shortest distance between the hand’s position and the predicted path of the ball. This creates a curve that shows how the hand approaches the ball over time. The shape of this curve reflects how the hand prepares for and moves toward the catch, giving insight into how the juggler plans and controls their movements. Looking at the process this way highlights how the hand actively works to meet the ball’s path, rather than just reacting to where the ball ends up.

We compared two juggling conditions, solo 3-ball cascade juggling (condition 1) and dyadic 3-ball cascade juggling (condition 2), with the independent variable being the availability of proprioceptive and tactile information from the throwing hand. Following the approach of Slupinski et al. (2018), we expected that jugglers can predict a ball’s trajectory earlier in solo juggling compared to dyadic juggling, and that the duration of the planned phase of the movement toward interception is longer in the solo case. This serves as our dependent variable. To our knowledge, this is the first study to examine both solo and dyadic juggling and he influence of internal information on the planning process during ball catching in toss juggling. Interestingly, in the dyadic scenario, the functional “deafferentation” leads to larger flight distances and, consequently, longer flight times. This aspect has been taken into account in the analyses presented later in the paper.

## 2 Materials and Methods

### 2.1 Participants

A total of 18 jugglers (12 males and 6 females) participated in the experiment. The jugglers’ age were in the range 20 - 60 years (mean = 30 years, SD = 10.42 years). The jugglers were chosen based on the inclusion criteria that they are able to comfortably sustain a 3 ball cascade pattern for longer than 20 s. All participants were either ambidextrous or right-handed. Handedness was assessed based on the participants’ preferred hand for daily activities and writing.

The participants received a monetary compensation of 8 € per hour or course credits, whichever they chose. The study was conducted following the Declaration of Helsinki 1964(Association 2013). The protocol was approved by the Ethical Review Board of the Justus Liebig University, Giessen, and subjects gave written informed consent to participate in the study.

### 2.2 Task and Setup

The experiment had two conditions. In the first condition, the juggler had to perform a regular 3 ball cascade pattern (solo juggling, Fig. 1a), and in the second condition, the juggler was paired with another juggler with whom they performed a side-to-side shared 3 ball cascade pattern (dyadic juggling, Fig. 1b). For the second pattern, both the jugglers stood next to each other facing forward, and the juggler (partner) standing to the left of the other juggler used their left hand, while the other juggler (initiator) used their right hand. After each trial, the jugglers switched places, i.e., the initiator of the previous trial became the partner in the next trial, and vice-versa. In both the conditions, each trial started with the first throw from the right hand (specifically, in the dyadic juggling case, by the initiator).

**Fig. 1.**
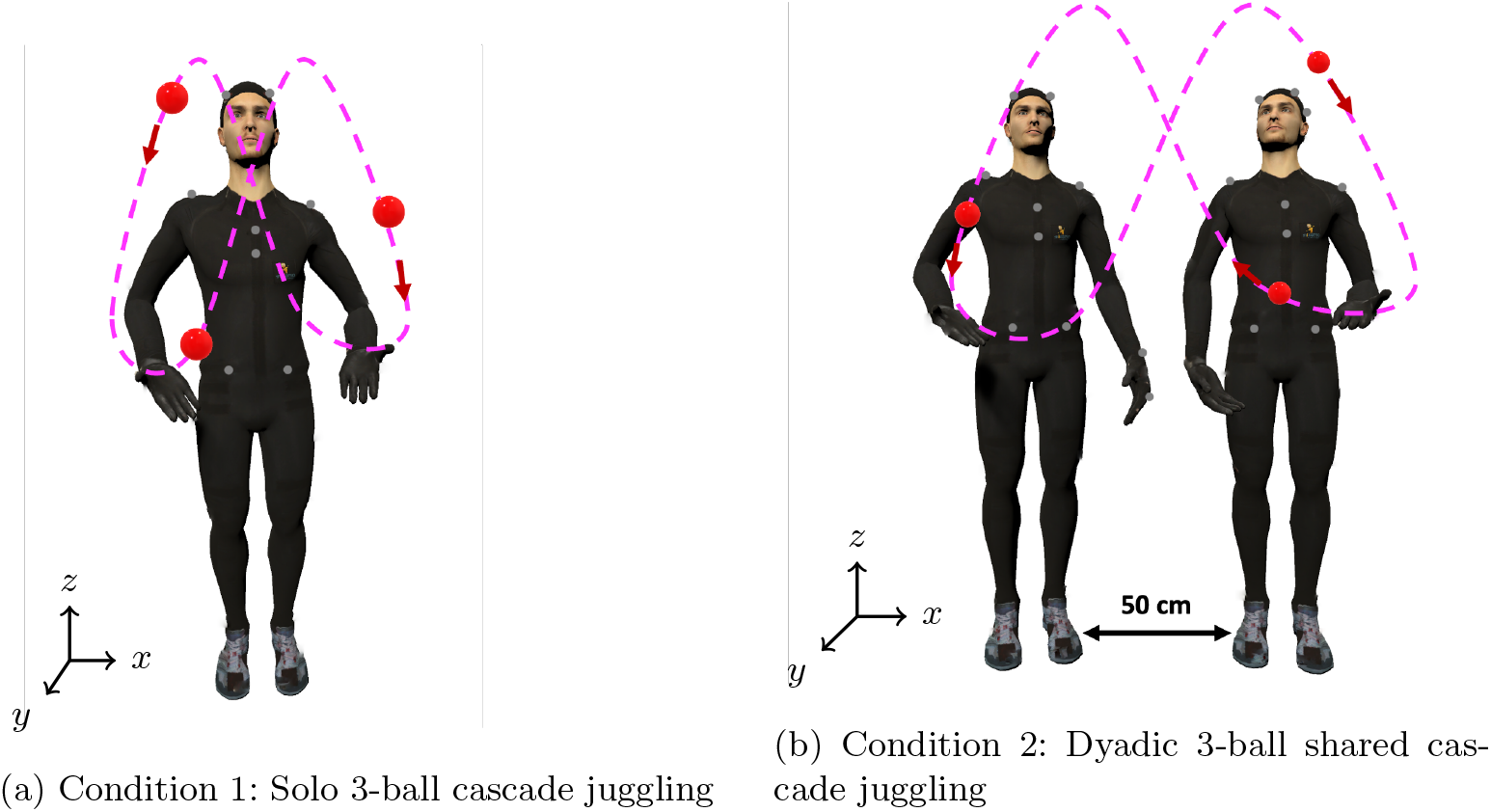
The juggling task involved in the two conditions of the experiment

No instructions regarding juggling speed and height were given, and the participants were instructed to start juggling after they heard the phrase “GO!” by the experimenter. In each condition, the participant had to complete 10 trials. Each trial lasted until the experimenter gave a “STOP!” command at around 20 seconds or until the juggling pattern collapsed, whichever happened first. Every trial started with the jugglers standing at the same spot and, in condition 2, they stood ≈ 50 cm apart (as illustrated in Fig. 1b).

### 2.3 Data Acquisition

Movements of the jugglers and the ball trajectories were recorded using a passive marker-based optoelectrical camera system (Vicon Motion Systems, Oxford, UK). 36 cameras (28 Vicon Vantage V5, and 8 Vicon Vero) were used and the data was collected at 240 Hz. The system was calibrated according to the manufacturer’s guidelines prior to data collection. Participants were taped with Pearl hard-base markers according to the upper body Plug-in-Gait marker set. Henrys smooth beanbag-style balls with ball diameters as 67 mm and weight 125 g were used as juggling balls. The three balls were covered with retro-reflective tape and defined as markers, to record the movements of the balls. The 3D trajectories of the passive markers were reconstructed using Nexus software (Vicon). The coordinate system used was defined as follows: 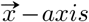 pointing left-to-right, 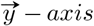 along the antero-posterior axis, and 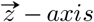 along the vertical axis. Additionally, the positive is upwards and the negative direction of the 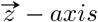 direction is downwards.

### 2.4 Data Processing and Analysis

Data processing and analyses were conducted using MATLAB R2024b (The Math-Works, Inc). The data was first filtered lightly with a first-order two-way (zero-lag) Butterworth low-pass filter with a 22 Hz cutoff frequency to smoothen out the raw motion capture data and remove the high frequency jitters. Next, the individual ball trajectories during flight time (i.e., the time between a ball’s release from one hand and its subsequent catch by the other) were segmented from looking at the vertical velocity profile of the balls to identify the moments of throw and catch. A positive velocity peak denotes the moment of throw, and a negative velocity peak denotes the moment of catch, as illustrated in Fig. 2. In the same duration, the trajectory of the markers placed on the dorsal head of the 3^*rd*^ metacarpal (FIN), and on the two sides of the wrist (WRA, WRB: thumb-side and “little finger”-side) were also segmented. In order to have a better representation of the hand position, which considers the offset between the hand and ball markers at contact, we did the following corrections. First, we transformed the ball-hand contact points to a hand coordinate system, taking the 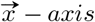 pointing left-to-right, the 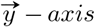 pointing back-to-front, and the 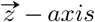 pointing upwards when the palm is facing upwards. The FIN marker was used as origin of the hand coordinate system (Fig. 3). This transformation of a global to a hand coordinate system of the hand-ball contact points enables detecting ball position relative to the hand. We then took the mean ball-contact position in this hand coordinate system separately for each hand and used this information to correct for the systematic offsets in ball-hand contact. This also included the accurate hand orientation for each frame.

**Fig. 2.**
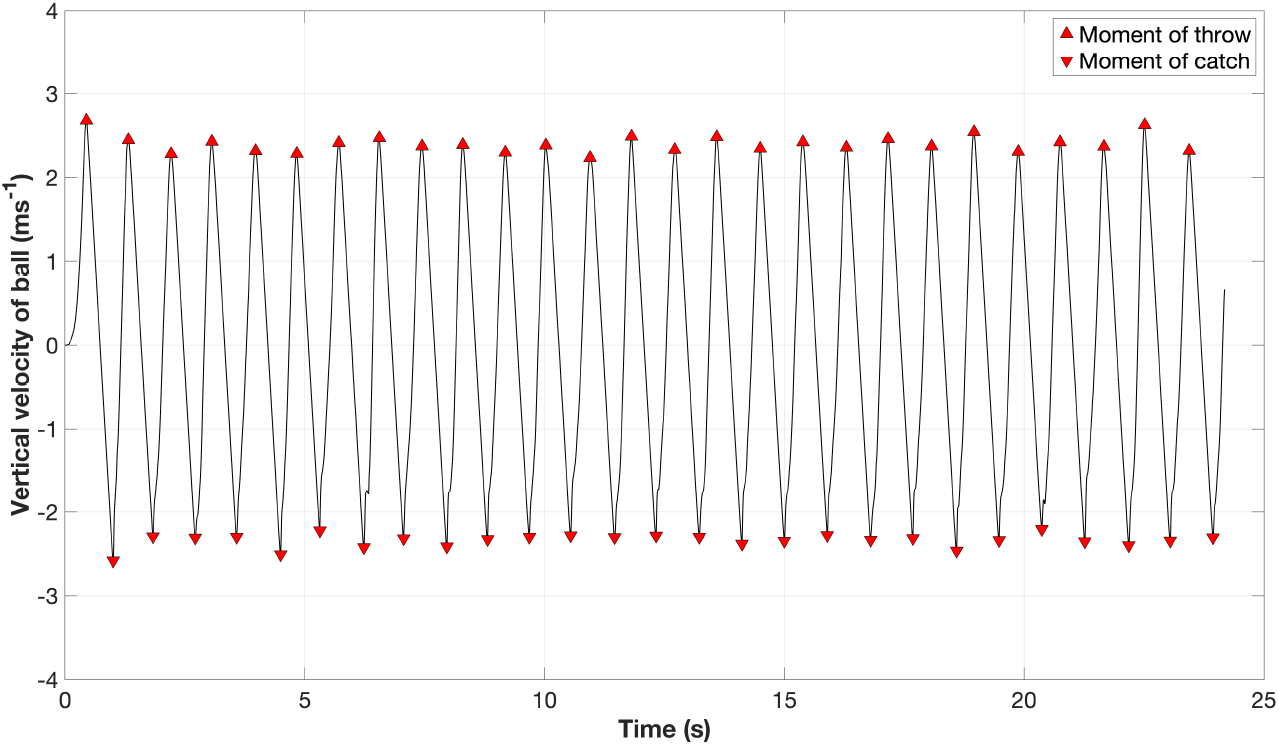
Vertical velocity profile of an example ball trajectory. The positive velocity peak corresponds to the ball release (throw), while the negative velocity peak corresponds to the ball interception (catch). The interval between two consecutive throw and catch markers represents the flight phase of the ball

**Fig. 3.**
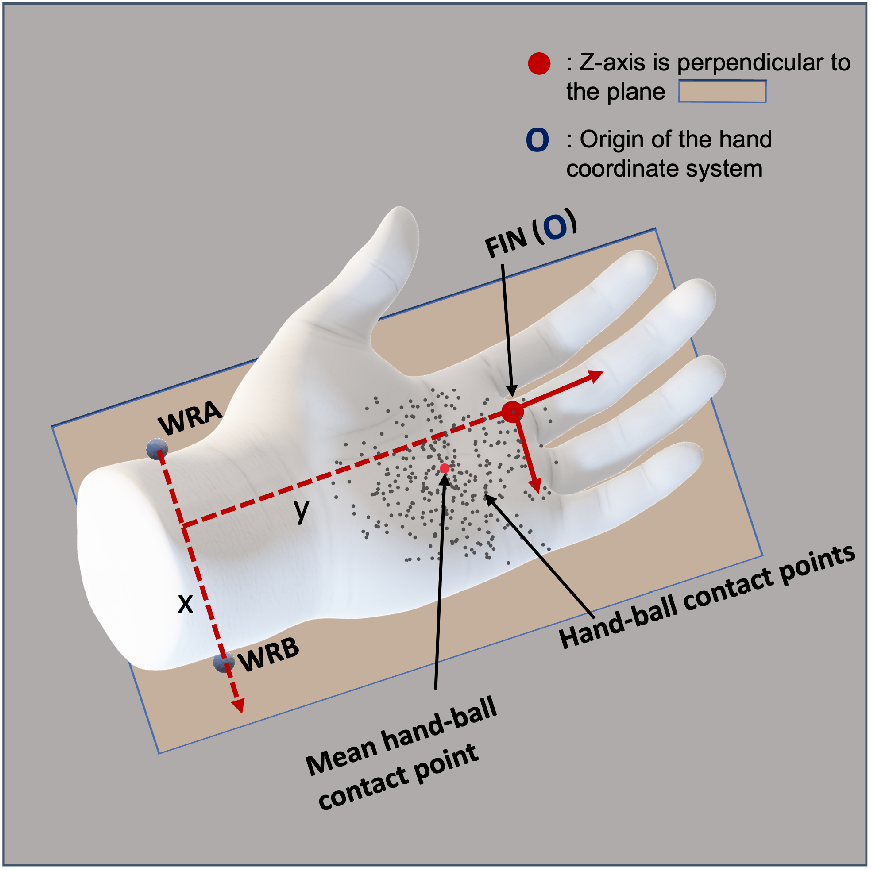
The hand representation in the analysis with 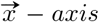 pointing left-to-right, the 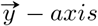 pointing back-to-front, and the 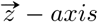 pointing up when the palm is facing upwards. The origin of this representation is at the FIN marker

This study explores to which extent anticipation of a ball’s trajectory based on internally available information during the throw affects the following catch of that particular ball. Following the method proposed in Slupinski et al. (2018), we examined how predictions based on information from the throwing hand affects the timing of the catching hand’s movements. By leveraging the empirical principle that goaldirected trajectories exhibit minimal jerk (Flash and Hogan 1985; Hogan 1984; Hoff 1992; Fligge et al. 2012; Desmurget et al. 1998), we investigated how early jugglers achieve such smooth, goal-directed hand movements in solo juggling (condition 1) compared to when they lack throwing information from the opposite hand (condition 2). This approach evaluates the shortest distance between the hand and the ball’s future trajectory over time. Using this one-dimensional trajectory data (MDHP or Minimum Distance of the Hand to the Parabola), the smooth (goal-directed) phase is determined by fitting it to the minimum jerk equation (Fig. 4), where jerk is defined as the third derivative of position with respect to time. A minimum jerk trajectory, *J*_*min*_, minimizes the sum of the squared jerk values along the object’s path between an initial time *t*_*o*_ and a final time *t*_*f*_ :

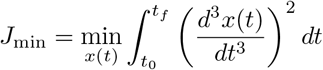

**Fig. 4.**
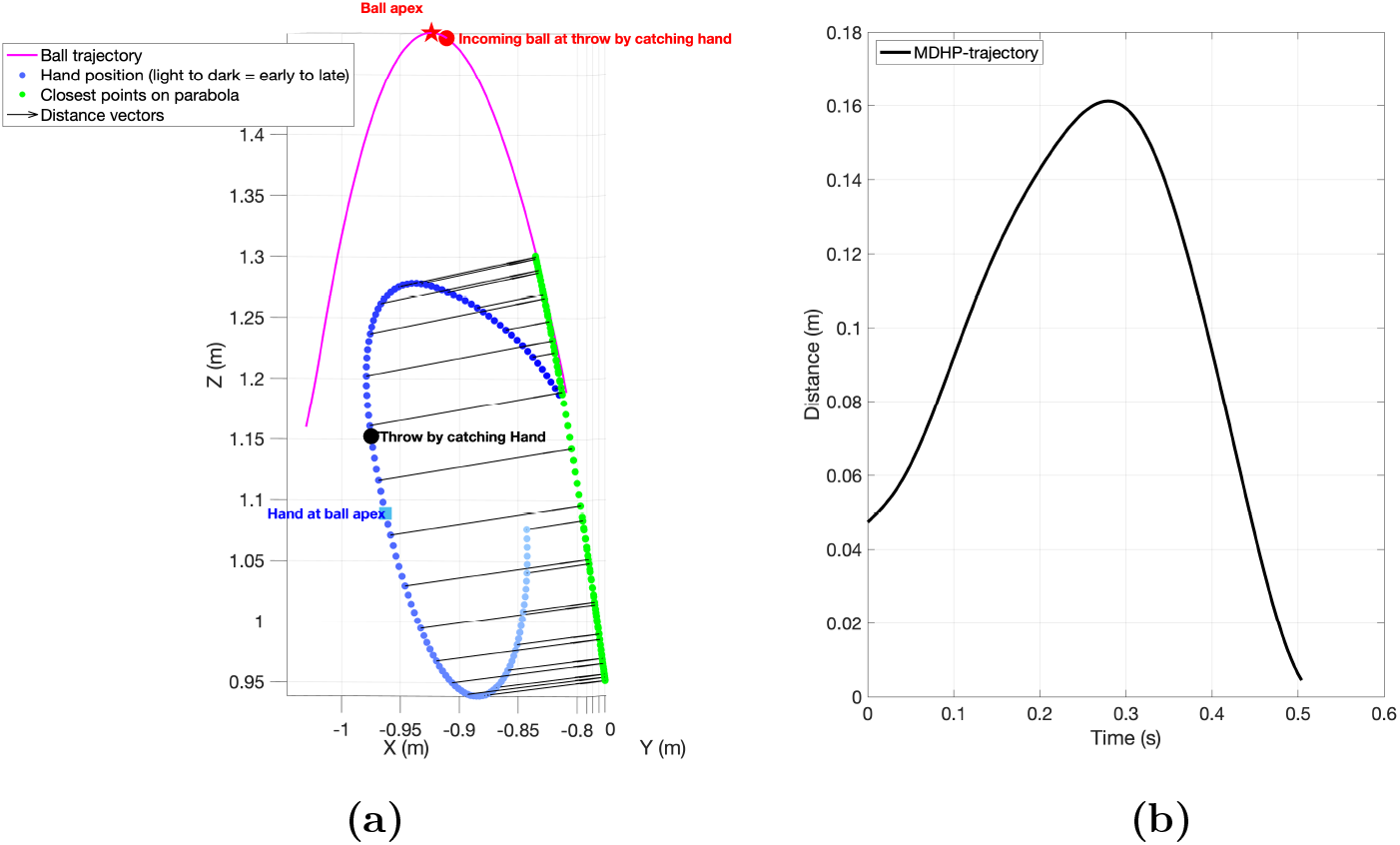
The figure in panel **(a)**. demonstrates the evolution of hand and ball trajectory, for an example trajectory, in the duration between a ball’s release from one hand and its subsequent catch by the other. The arrows mark the minimum distance vectors from hand to ball’s future trajectory, plotted every 5 time samples apart (to avoid visualization clutter). The figure in panel **(b)**. demonstrates the corresponding Minimum Distance of the Hand to the Parabola (MDHP) trajectory for the same example trajectory

To prepare for analysis, we first filtered the MDHP trajectory with a third-order (zero-lag) Butterworth low-pass filter with a 12 Hz cutoff frequency. We next segment the MDHP trajectory into its constituent functional phases for analysis. The first phase in the segmentation constitutes the approximation to the parabola phase and the next phase is the interception along the parabola phase. The first approximation phase starts from the time when the ball leaves the other hand and ends at the end of the goal-directed movement phase. The next phase marks the start of the interception phase, where the hand could move along the trajectory of the ball and make the catch. This time point *t*_*endSmoothApprox*_ is the time when the hand is still approaching the ball’s parabola but very slowly. The catcher’s hand was considered to follow the ball’s trajectory closely enough for a successful catch when the Minimum Distance of the Hand to the Parabola (MDHP) drops below 0.07 m before the actual interception of the ball. In order to ensure that the search for *t*_*endSmoothApprox*_ happens after the ball already in the catching hand has been released, we choose the starting time of the window as either the moment the ball already in hand was thrown or the point when the MDHP first dropped below 0.07 m, whichever happened later. Eventually we identified the first peak deceleration in the MDHP trajectory as the start of the interception phase (*t*_*endSmoothApprox*_), as illustrated in Fig. 5.

**Fig. 5.**
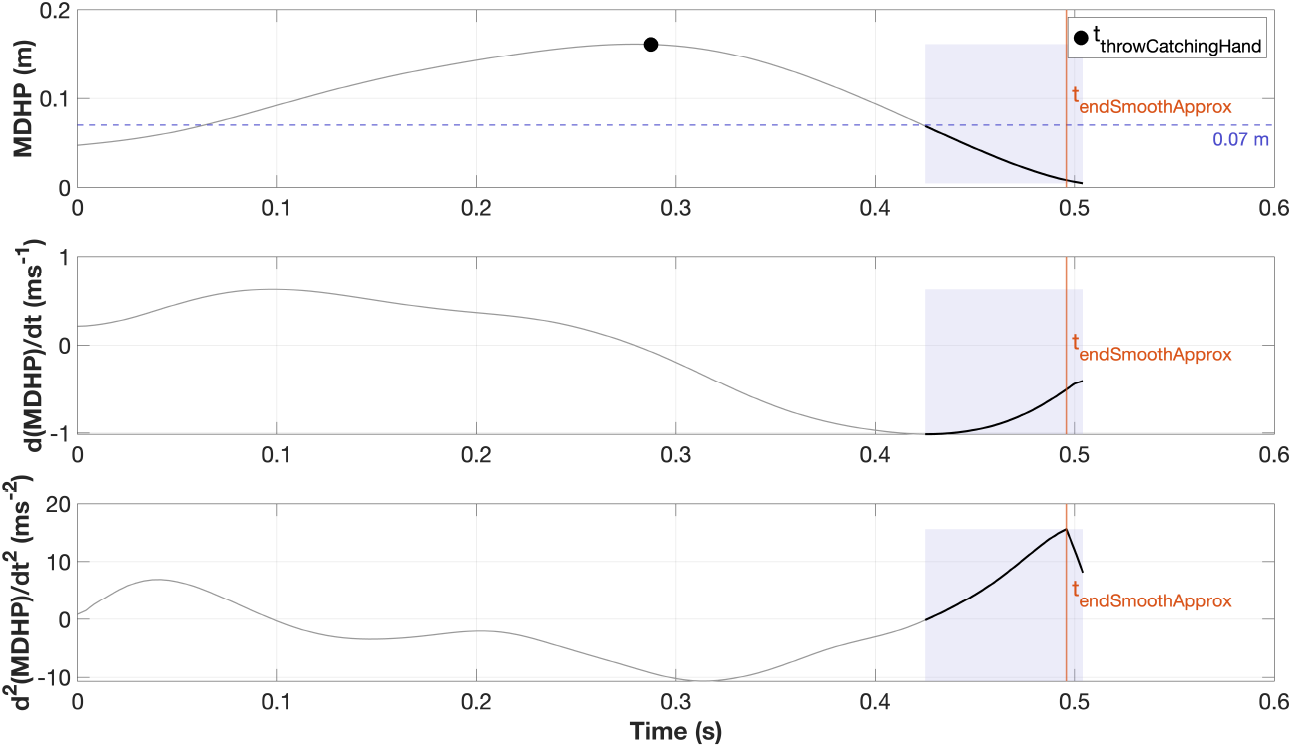
Identification of *t* _*endSmoothApprox*_ in the MDHP trajectory. MDHP versus time with shaded region where MDHP *<* 0.07 m, threshold line at 0.07 m (blue dashed), and marker indicating the throw by the catching hand (black filled circle). The vertical line marks *t*_*endSmoothApprox*_, defined as the first peak deceleration in the MDHP trajectory, obtained by looking at the second derivative of MDHP trajectory and identifying the first peak within the shaded region

### 2.5 Dependent Variable

The dependent variable, the start of the smooth approximation phase, was determined following the method described in Slupinski et al. (2018). The variable, (*t*_*startSmoothApprox*_), denotes the time the planning for the arm movement is concluded and consequently reflects the catcher’s anticipation of the ball trajectory. It is the earliest time at which the MDHP-trajectory satisfied the minimum jerk criterion (Flash and Hogan 1985; Hogan 1984). The minimum jerk trajectories for each time step before *t*_*endSmoothApprox*_ were calculated until the root mean square error (RMSE) in position exceeded 1 mm, considering the spatial resolution of a general motion capture system (Slupinski et al. 2018) (as illustrated in Fig. 6).

**Fig. 6.**
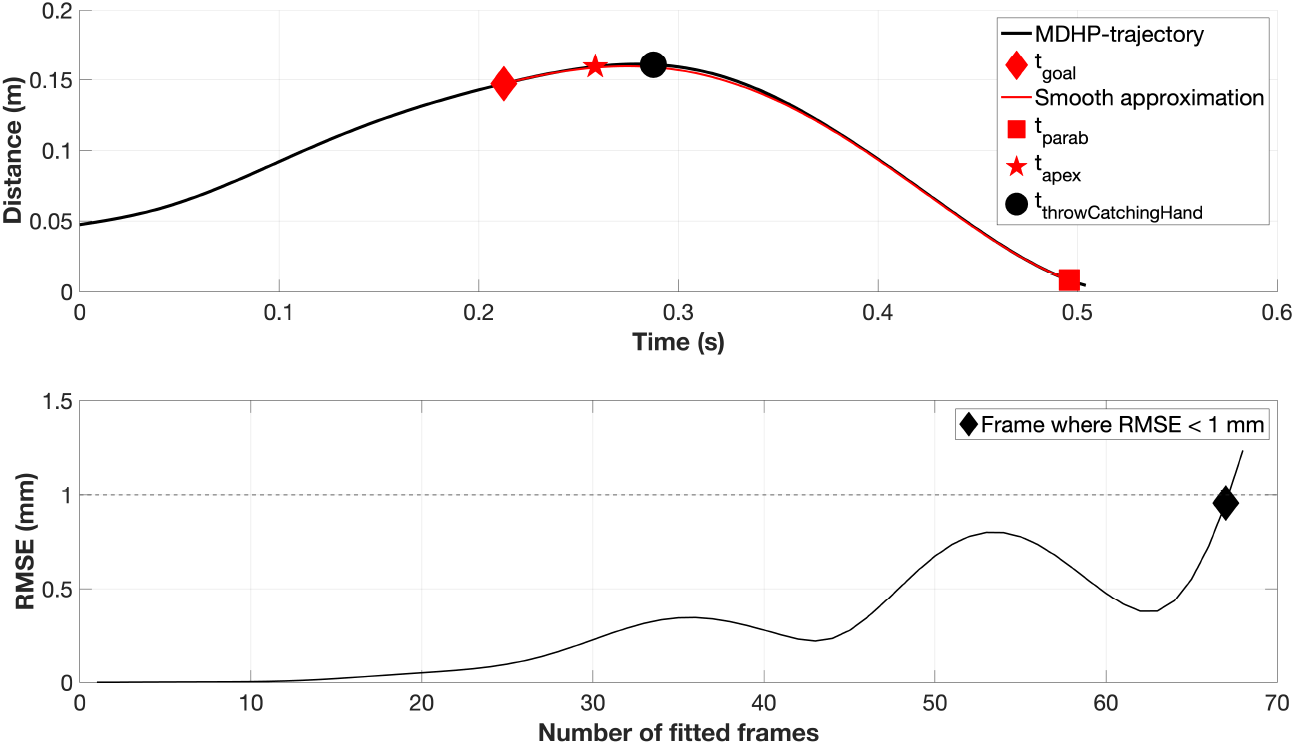
Example of a minimum jerk fit on the MDHP trajectory for a solo condition. Minimum jerk trajectories are plotted for each time step before *t*_*endSmoothApprox*_ and continue until the RMSE of the fit exceeds 1 mm, which marks the end of the fit. The resulting minimum jerk trajectory is the smooth approximation curve (as illustrated), which functionally starts from *t*_*startSmoothApprox*_ and ends at *t*_*endSmoothApprox*_. In addition, *t*_*apex*_ and *t*_*throwCatchingHand*_ denote the moment when the incoming ball reaches its apex and the moment when the catching hand is emptied before intercepting the incoming ball, respectively

To meaningfully eliminate variability by enabling comparison in flight times and, subsequently, the relevant temporal variables in our analysis, we have normalized time to the ball’s apex and the throw of ball by the catching hand, respectively. The apex was chosen as the anchor point under the assumption that jugglers use the zenith of the ball’s flight as a key reference for their prediction, since this is where the ball’s vertical velocity is zero and trajectory information is clustered there (Van Santvoord and Beek 1994; Amazeen et al. 1999; Huys and Beek 2002; Dessing et al. 2012; Beek and Lewbel 1995). Additionally, absolute time, governed by the external clock, reflects the real world temporal evolution of each event. However, the motor control processes underlying skilled actions such as juggling are often organized relative to internal timing mechanisms that operate independently of physical time. By normalizing each throw’s time axis, we map the movement data onto a common internal timescale, capturing the phase progression of each action. This also allows us to compare the onset of the smooth approximation phase with different events within the trajectory, ensuring that the event in question is temporally aligned at the same point for both conditions. In this way, we can examine whether the hand’s approach to the ball is consistently structured with respect to the internal dynamics of an event in the juggling cycle, rather than tied to specific points on the external timeline.

All parameters refer to time (in s) within a single trajectory with *trajDuration* being the duration of the trajectory, and each measure is expressed as a percentage of the remaining flight duration from the reference point to the interception.

The timing of the outgoing throw relative to the apex of the incoming ball’s flight was defined as:

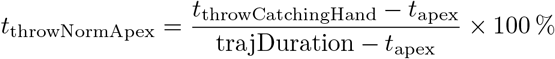

The onset of the smooth approximation phase (*t*_startSmoothApprox_) relative to the apex was defined as:

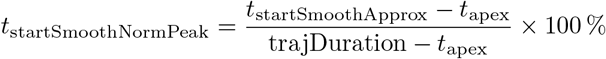

The timing of the apex relative to the outgoing throw that frees the catching hand was computed as:

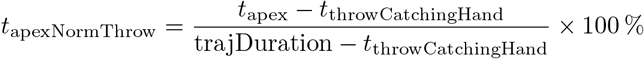

Finally, the onset of the smooth approximation phase relative to the outgoing throw was calculated as:

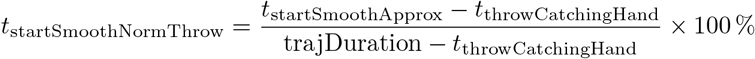

### 2.6 Statistical Analysis

JASP version 0.19.3 (JASP Team 2025) was used for all statistical analyses. For all tests, the significance level was set to *α* = 0.05. Due to skewness in the individual participant data, median for all parameters were calculated for each participant. The necessary normality assumption to run a paired *t* -test was checked using the Shapiro-Wilk test, which indicated no significant deviations from normality. Finally, a one-tailed paired *t* -test was used to compare the dependent variables between two conditions. Statistical effect sizes were determined using Cohen’s *d* measure and they were corrected for correlation between observations (Lakens 2013).

A priori power analysis (G^*^Power) was conducted with a conservative effect size of 0.8 (following Slupinski et al. (2018) but allowing extra variability due to the juggling task), *α* = 0.05, and power = 0.95, yielding a required sample size of 18 participants.

## 3 Results

### 3.1 Start of Smooth Approximation

Fig. 7 shows the distribution of onset timing of smooth approximation phase (*t*_*startSmoothApprox*_) in absolute time. It is evident from the figure that, in condition 1, *t*_*startSmoothApprox*_ started significantly earlier compared to condition 2 [*t*_17_ = −8.251, *p<* 0.001, Cohen’s *d* =−1.990]. Interestingly, the onset of *t*_*startSmoothApprox*_ occurred more frequently before the incoming ball’s apex in condition 1 than in condition 2.

**Fig. 7.**
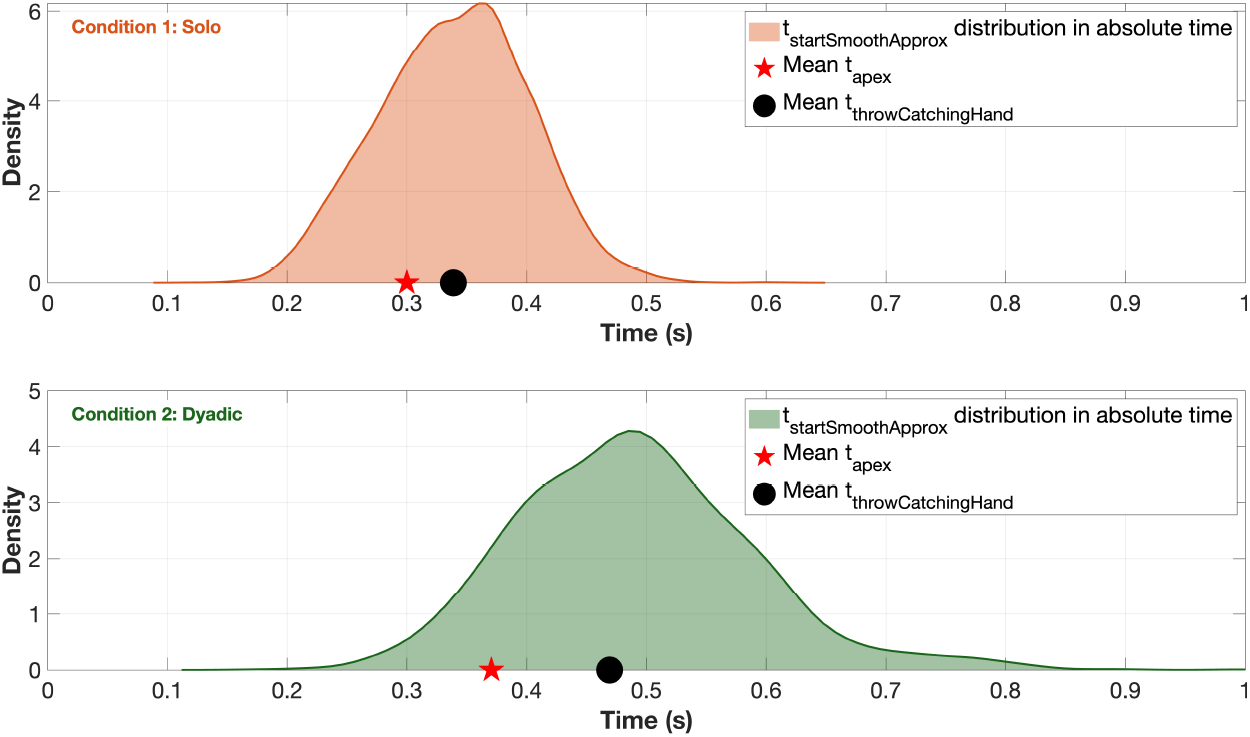
Distribution of *t*_*startSmoothApprox*_ for condition 1 & 2 in non-normalized time. Red markers indicate the mean ball apex times, and black markers indicate the mean times of the ball throws preceding interception by the intercepting hand

Since ball flight times varied across trials, aligning movements to a common internal timescale allows direct comparison of event timing across conditions. In Fig. 8, we visualize the normalized timing of smooth approximation onset (*t*_*startSmoothNormP eak*_) with respect to the incoming ball’s apex and expectedly observe that the start of the smooth approximation phase with respect to apex was significantly earlier [*t*_17_ = −10.905, *p<* 0.001, Cohen’s *d* =−2.095] in the solo juggling scenario compared to its dyadic counterpart. Here, 0% corresponds to the moment when the incoming ball reaches its apex, and 100% marks the actual interception. To account for the time required to initiate corrective movements, we included an additional window of 110 ms, based on established findings in interception research (Brenner et al. 1998; Soechting and Lacquaniti 1983; Brenner and Smeets 1997; Fooken et al. 2024), which show that when a moving target is visually perturbed, the hand’s corrective response latency is approximately 110 ms. In Fig. 8, this window is time-normalized relative to the average trajectory length for each condition and aligned to the apex, because if jugglers were merely reacting to the ball apex and updated their plan at that moment, it would have taken at least 110 ms for the corrective movement. In contrast, it can be observed that the onsets of the smooth-approximation phase fall mostly into this window in the solo condition compared to the dyadic condition.

**Fig. 8.**
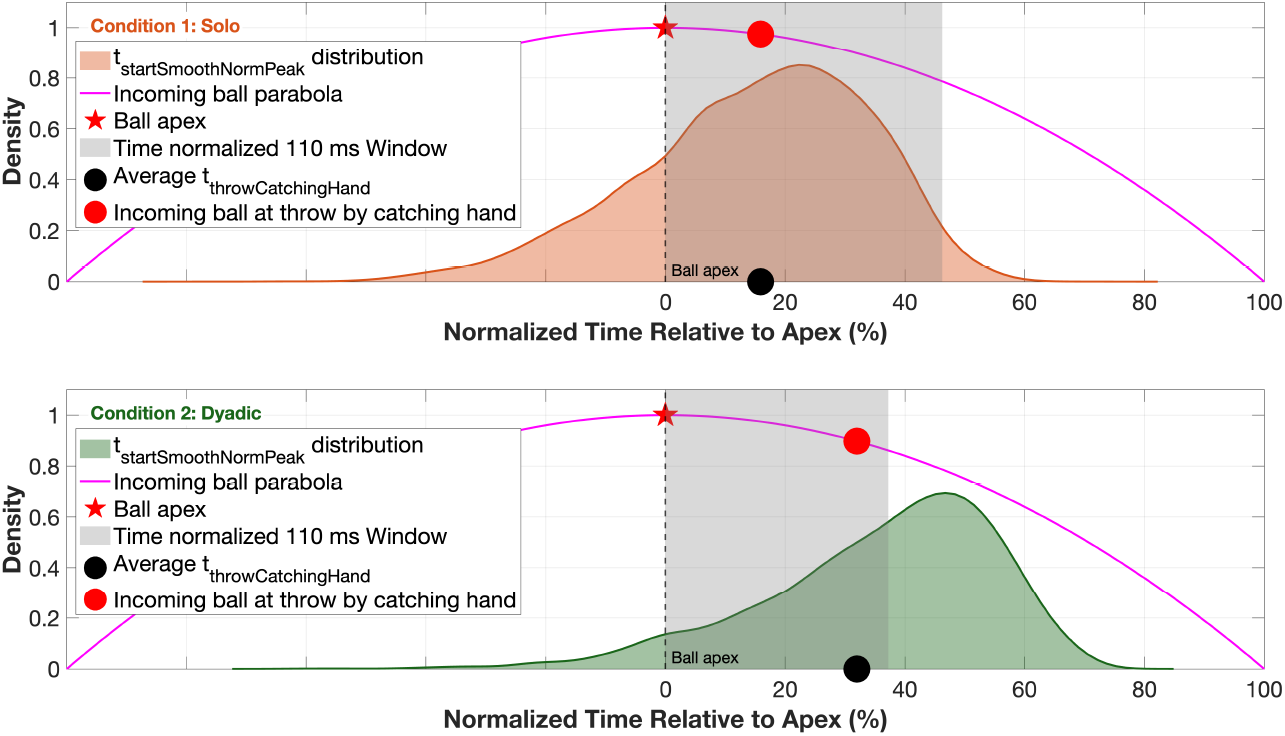
Distribution of normalized onset timing of smooth approximation phase relative to the incoming ball’s apex, *t*_*startSmoothNormP eak*_, for condition 1 & 2. 0 % indicates the event of the ball reaching its apex and 100% marks the interception. The shaded area represents a 110 ms visuomotor response window, adjusted to the average flight duration for each condition. Condition 1 shows an earlier onset of the smooth approximation phase relative to the apex compared to condition 2. The incoming ball parabola is plotted for illustration purposes, indicating different events in the trajectory, i.e., the ball apex and the incoming ball’s position at the throw by the catching hand

### 3.2 Testing Competing Hypotheses

In both conditions, jugglers make an outgoing throw to empty the catching hand for the next catch. If the preparation time for the catching action were always the same after freeing the hand, no difference would be expected between conditions. However, Fig. 9 illustrates that the timing of the onset of the smooth approximation phase (*t*_*startSmoothNormT hrow*_) cannot be explained simply by when the catching hand is freed. The onset of the smooth phase occurs significantly later [*t*_17_ = −3.448, *p<* 0.05, Cohen’s *d* = −1.174] relative to the outgoing throw in the dyadic condition compared to the solo condition.

**Fig. 9.**
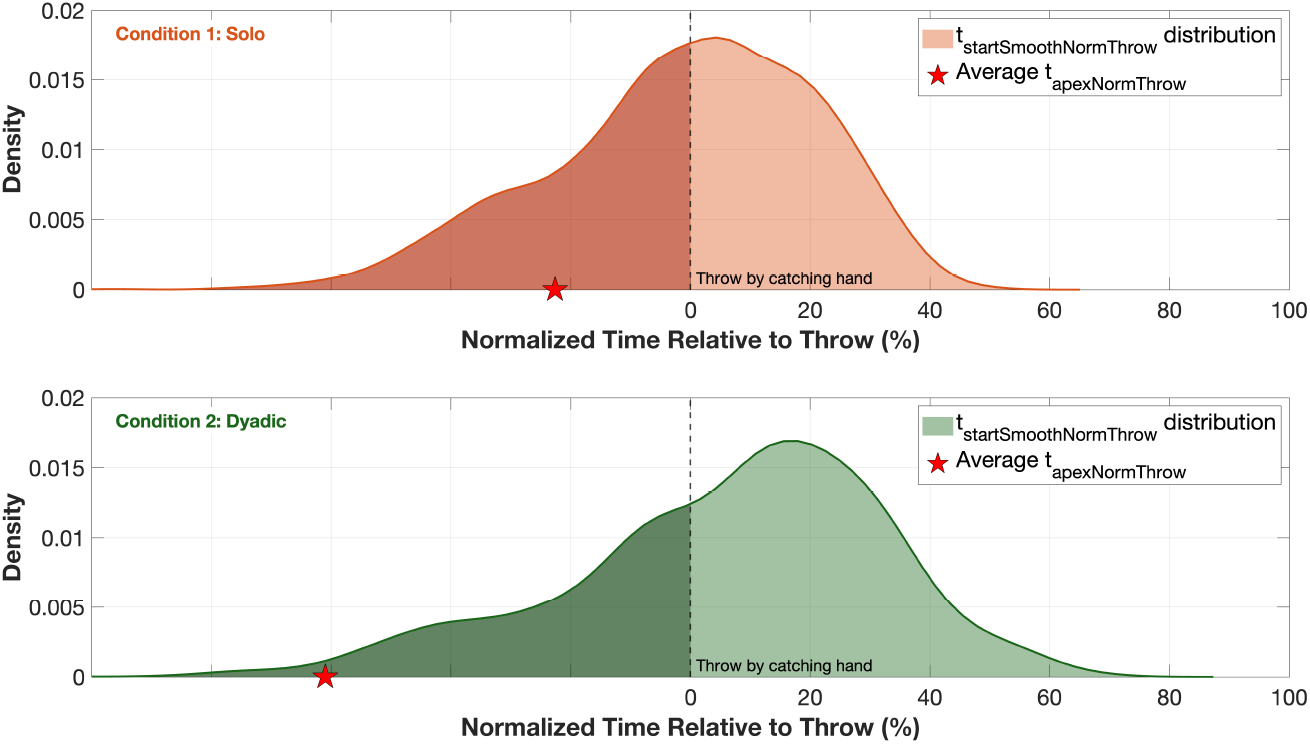
Distribution of normalized onset timing of smooth approximation phase defined relative to the ball throw preceding interception by the intercepting hand, *t*_*startSmoothNormT hrow*_. The average normalized ball apex relative to the outgoing throw that frees the catching hand, *t*_*apexNormT hrow*_, for each condition is also marked with a red pentagram. The proportions of the shaded regions preceding the throw by the catching hand can be visually compared between the two conditions

Additionally, the duration of the smooth approximation phase, defined as the interval from *t*_*startSmoothApprox*_ to *t*_*endSmoothApprox*_, was analyzed to test whether longer absolute flight times in the dyadic condition could account for the delayed onset observed previously. Fig. 10 shows that the duration of the smooth phase was significantly shorter [*t*_17_ = 5.563, *p<* 0.001, Cohen’s *d* =1.358] in the dyadic condition compared to the solo condition.

**Fig. 10.**
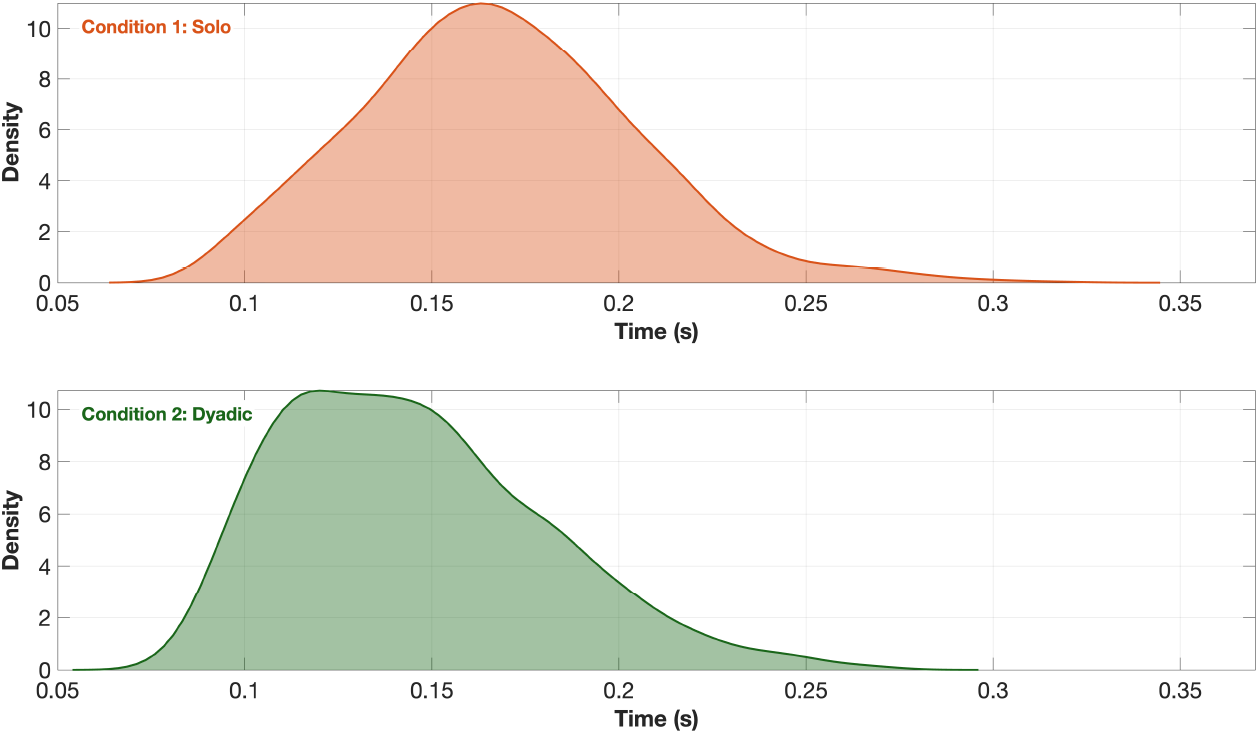
Distribution of smooth approximation durations, measured as the time between *t*_*startSmoothApprox*_ and *t*_*endSmoothApprox*_ (s)

## 4 Discussion

In this study, we wanted to investigate the relevance of internal and tactile information from one’s own throwing movement for the planning of anticipatory movement parts of a subsequent catch. To this end, we employed a paradigm that included both self-generated actions (solo juggling) and externally generated actions (dyadic juggling). The dyadic condition is our best attempt to “deafferent” a person in order to study interception during juggling. Our specific aim was to investigate whether the absence of internal information about the throw, such as when the throw is made by a partner, affects the timing of hand movement planning for the subsequent catch. The answer to this question is not particularly straightforward in this context, since planning runs parallel with ongoing movement control. The juggler must release the ball currently held in the hand before initiating the catch movement, all while maintaining the temporal stability of the juggling rhythm. Our underlying assumption was that anticipatory behavior would be reflected in the timing at which the hand smoothly moves toward the trajectory of the incoming ball. Our findings align with prior work showing earlier planning conclusion and movement initiation for self-generated discrete throws (Hagenfeld et al. 2022; Slupinski et al. 2018), but critically, we demonstrate this effect in a continuous coordinated task. This suggests that predictive control strategies are not limited to discrete reaches, but are continuously adapted based on the internal vs. external origin of motor timing information, even in non-discrete, continuous tasks like toss juggling.

Unlike Slupinski et al. (2018) and Hagenfeld et al. (2022), we could not fully control for ball trajectory flight time and flight distance due to the dynamic and rhythmic nature of the juggling task. However, we accounted for this variability in our analysis by normalizing times to flight duration. One could argue that this variation in flight time provided an opportunity to test whether having more time available leads to earlier movement onset. Since the hand is emptied in both conditions, following the alternative explanation, the preparation time should be similar if freeing the hand was the only constraint. Following this explanation, we should not see a significant shift in the onset of smooth phase distributions between the two conditions. Fig. 9 shows otherwise and rules out the alternative explanation that the timing difference in *t*_*startSmoothApprox*_ is solely due to when the hand is freed for the catch. The delay seen in the dyadic condition is likely due to the absence of internal motor information. When jugglers throw the ball themselves, as in the solo condition, they have access to efferent and proprioceptive signals that help them plan the catch. In the dyadic condition, this information is missing, which likely delays the initiation of predictive control. Additionally, if the timing difference was only a result of having more time as in condition 2, then the duration of the smooth, goal-directed phase should be similar or longer. However, Fig. 10 rejects the alternative explanation that the delayed onset of the smooth phase in dyadic juggling is solely due to reduced time pressure from longer ball flights. The shorter smooth phase indicates that jugglers delayed committing to predictive control even when more time was available. This supports the interpretation that the absence of efferent and proprioceptive information from self-generated throws diminishes the timely initiation of prediction in the dyadic condition. Interestingly, this reflects broader findings that relying only on external sensory motion cues can limit predictive accuracy. For instance, Kreyenmeier et al. (2022) showed that when people had to predict the reappearance of an accelerating target using visual information alone, they systematically mistimed their interception once the target was briefly occluded. Taken together, our results indicate that it is not simply the amount of time available that limits movement planning, but rather the level of predictive certainty.

It may seem surprising that adjustments to movement can occur right at the start, given the physical distance between the brain and the arm muscles, as well as the inherent physiological delay in muscle activation. Although responses to visual disturbances can be initiated in as little as ∼110 milliseconds (Brenner et al. 1998; Brenner and Smeets 1997), this is still relatively slow compared to the total duration of a catching movement. In fact, even movements traditionally considered ballistic are under continuous visual control. Additionally, the acceleration of the hand also modulates in real-time in response to target speed changes, with effects visible after roughly 200 ms (Brenner et al. 1998). The apex of the ball’s trajectory was chosen as an anchor point for analyzing the onset of predictive control because it provides an ideal moment for jugglers to update their internal estimates about the ball’s flight. At the apex, the ball’s vertical velocity is momentarily zero, making its position and path maximally stable and predictable. Prior studies have shown that jugglers naturally rely on visual information near the apex; novices often directly fixate on each ball’s apex to judge its landing point, while experienced jugglers implicitly use apex cues to maintain timing and consistency (Van Santvoord and Beek 1994; Huys and Beek 2002; Dessing et al. 2012; Amazeen et al. 1999). Even when vision is restricted to a narrow region near the apex, skilled jugglers can maintain the pattern, demonstrating how the apex contains key predictive information about trajectory and timing. This is directly relevant to how we interpret the 110 ms window shown in Fig. 8. If jugglers were relying only on this apex information to adjust their movement, any correction triggered around this point would require at least about 110 ms to appear in the hand’s motion, given typical visual processing and neuromuscular delays (Brenner et al. 1998; Brenner and Smeets 1997). In juggling, this dynamic plays out clearly as the apex may act as a reliable visual reference for feedforward prediction, but jugglers can also make fine-tuned corrections in the final phase before interception. The 110 ms window reflects this limit, any visual corrections initiated after this point are unlikely to influence the outcome in time. Thus, when the onset of the smooth approximation phase (*t*_*startSmoothApprox*_) falls predominantly within this window, as seen in the solo condition, it suggests that jugglers are relying more on preplanned, internally generated predictions. In contrast, when *t*_*startSmoothApprox*_ shifts later, as in the dyadic condition, this indicates increased dependence on continuous visual feedback to refine the interception trajectory. This distinction aligns with the broader assumption that skilled juggling combines both feedforward and feedback control, supported by efference copies of outgoing motor commands and internal forward models (Miall and Wolpert 1996) that help predict the consequences of the movement in real time.

To finally summarize, the goal of this study was to demonstrate that the internal information from the juggler’s throwing hand plays a critical role in determining the timing of the catching hand’s movement. In a juggling task, where the ball can be caught at any point along its parabolic trajectory, accurate and early prediction of the ball’s path is essential for maintaining stability in the juggling pattern. We found that jugglers use information from their own throw (when available) to make those predictions early, enabling a timely and smooth interception. Our findings highlight the relationship between sensory input and motor planning, emphasizing the importance of efferent, proprioceptive, and tactile information in enabling smoother and more efficient hand movements in complex sensorimotor tasks like juggling. The juggling example further illustrates how the brain brings together feedforward and feedback control. Feedforward predictions based on efference copy and internal forward model allows the catching movement to be initiated early and in the right general direction, while feedback (mainly visual, especially near the apex or during ball flight) provides ongoing corrections to fine tune the movement, which is under control of the perceived position of the target. When internal information are available (as in solo juggling), the system relies more on predictive control, when they are absent (as in catching someone else’s throw in dyadic juggling), the system increases reliance on prospective control. Ultimately, jugglers achieve a seamless integration of these processes. They demonstrate that even a task often perceived as rapid and rhythmic still involves guidance by sensory input, be it the “internal” vision of one’s own throw or the “external” sight of the ball in flight. This synergy between sensory input and motor planning is fundamental not only to juggling, but to a wide range of interceptive and interactive motor behaviors in daily life.

## 5 Declarations

### Author Contributions

All authors contributed to the study conception and design. Abir Chowdhury and Heiko Maurer contributed to the experimental setup. Abir Chowdhury performed the data collection and formal data analysis. Abir Chowdhury and Heiko Maurer contributed to the software. Abir Chowdhury drafted the first version of the manuscript. All authors contributed to the interpretation of results, provided critical feedback on manuscript revisions, and approved the final version of the manuscript.

## Acknowledgements

We would like to thank Philipp Ewen for his assistance with data collection.

## Data Availability

Data used in this study will be made public at osf.io/u8b3r upon acceptance for publication.

## Informed Consent

All participants gave written informed consent.

## Ethics Approval

The study complied with the approval by the Ethical Review Board of the Justus Liebig University, Giessen.

## References

Amazeen EL, Amazeen PG, Post AA, Beek PJ (1999) Timing the selection of information during rhythmic catching. Journal of Motor Behavior 31(3):279–289. 10.1080/00222899909600994

Association WM (2013) World medical association declaration of helsinki: ethical principles for medical research involving human subjects. Journal of the American Medical Association 310(20):2191–2194. 10.1001/jama.2013.281053

Beek PJ, Lewbel A (1995) The science of juggling. Scientific American 273(5):92–97

Beek PJ, Turvey MT (1992) Temporal patterning in cascade juggling. Journal of Experimental Psychology: Human Perception and Performance 18(4):934. 10.1037/0096-1523.18.4.934

Botvinick-Greenhouse J, Shinbrot T (2020) Juggling dynamics. Physics Today 73(2):62–63. 10.1063/PT.3.4417

Brenner E, Smeets JBJ (1997) Fast responses of the human hand to changes in target position. Journal of Motor Behavior 29(4):297–310. 10.1080/00222899709600017

Brenner E, Smeets JBJ, De Lussanet MH (1998) Hitting moving targets continuous control of the acceleration of the hand on the basis of the target’s velocity: Continuous control of the acceleration of the hand on the basis of the target’s velocity. Experimental Brain Research 122(4):467–474. 10.1007/s002210050535

Desmurget M, Pélisson D, Rossetti Y, Prablanc C (1998) From eye to hand: planning goal-directed movements. Neuroscience & Biobehavioral Reviews 22(6):761–788. 10.1016/s0149-7634(98)00004-9

Dessing JC, Oostwoud Wijdenes L, Peper CE, Beek PJ (2009) Visuomotor transformation for interception: catching while fixating. Experimental Brain Research 196:511–527. 10.1007/s00221-009-1882-6

Dessing JC, Rey FP, Beek PJ (2012) Gaze fixation improves the stability of expert juggling. Experimental Brain Research 216:635–644. 10.1007/s00221-011-2967-6

Diaz G, Cooper J, Rothkopf C, Hayhoe M (2013) Saccades to future ball location reveal memory-based prediction in a virtual-reality interception task. Journal of Vision 13(1):20–20. 10.1167/13.1.20

Flash T, Hogan N (1985) The coordination of arm movements: an experimentally confirmed mathematical model. Journal of Neuroscience 5(7):1688–1703. 10.1523/JNEUROSCI.05-07-01688.1985

Fligge N, McIntyre J, van der Smagt P (2012) Minimum jerk for human catching movements in 3D. In: 2012 4th IEEE RAS & EMBS International Conference on Biomedical Robotics and Biomechatronics (BioRob), IEEE, pp 581–586, 10.1109/BioRob.2012.6290265

Fooken J, Balalaie P, Park K, Flanagan JR, Scott SH (2024) Rapid eye and hand responses in an interception task are differentially modulated by context-dependent predictability. Journal of Vision 24(12):10–10. 10.1167/jov.24.12.10

Hagenfeld L, de Lussanet MHE, Boström KJ, Wagner H (2022) Planning catching movements: Advantages of expertise, visibility and self-throwing. Journal of Motor Behavior 54(5):548–557. 10.1080/00222895.2021.2022591

Hayhoe M, Ballard D (2005) Eye movements in natural behavior. Trends in Cognitive Sciences 9(4):188–194. 10.1016/j.tics.2005.02.009

Hoff B (1992) A model of the effects of speed, accuracy, and perturbation on visually guided reaching. Experimental Brain Research 22:285–306

Hogan N (1984) Adaptive control of mechanical impedance by coactivation of antagonist muscles. IEEE Transactions on Automatic Control 29(8):681–690. 10.1109/TAC.1984.1103644

Huys R, Beek PJ (2002) The coupling between point-of-gaze and ballmovements in three-ball cascade juggling: the effects of expertise, pattern and tempo. Journal of Sports Sciences 20(3):171–186. 10.1080/026404102317284745

JASP Team (2025) JASP (Version 0.19.3)[Computer software]. URL https://jasp-stats.org/

Kawato M (1999) Internal models for motor control and trajectory planning. Current Opinion in Neurobiology 9(6):718–727. 10.1016/S0959-4388(99)00028-8

Kreyenmeier P, Kämmer L, Fooken J, Spering M (2022) Humans can track but fail to predict accelerating objects. eNeuro 9(5). 10.1523/ENEURO.0185-22.2022

Lakens D (2013) Calculating and reporting effect sizes to facilitate cumulative science: a practical primer for t-tests and anovas. Frontiers in Psychology 4:863. 10.3389/fpsyg.2013.00863

López-Moliner J, Brenner E, Louw S, Smeets JBJ (2010) Catching a gently thrown ball. Experimental Brain Research 206:409–417. 10.1007/s00221-010-2421-1

de Lussanet MH, Smeets JBJ, Brenner E (2004) The quantitative use of velocity information in fast interception. Experimental Brain Research 157:181–196. 10.1007/s00221-004-1832-2

Miall RC, Wolpert DM (1996) Forward models for physiological motor control. Neural Networks 9(8):1265–1279. 10.1016/s0893-6080(96)00035-4

Shadmehr R, Wise SP (2004) The computational neurobiology of reaching and pointing: a foundation for motor learning. MIT press

Slupinski L, de Lussanet MHE, Wagner H (2018) Analyzing the kinematics of hand movements in catching tasks—an online correction analysis of movement toward the target’s trajectory. Behavior Research Methods 50(6):2316–2324. 10.3758/s13428-017-0995-2

Soechting JF, Lacquaniti F (1983) Modification of trajectory of a pointing movement in response to a change in target location. Journal of Neurophysiology 49(2):548– 564. 10.1152/jn.1983.49.2.548

Van Santvoord AAM, Beek PJ (1994) Phasing and the pickup of optical information in cascade juggling. Ecological Psychology 6(4):239–263. 10.1207/s15326969eco06041

Wolpert DM, Ghahramani Z, Jordan MI (1995) An internal model for sensorimotor integration. Science 269(5232):1880–1882. 10.1126/science.7569931

Wolpert DM, Miall RC, Kawato M (1998) Internal models in the cerebellum. Trends in Cognitive Sciences 2(9):338–347. 10.1016/S1364-6613(98)01221-2

